# Decoding stimulus-specific regulation of promoter activity of p53 target genes

**DOI:** 10.1101/2025.04.09.647556

**Authors:** Flavia Vigliotti, Nicolai Engelmann, Mark Sinzger-D’Angelo, Heinz Koeppl, Alexander Loewer

**Affiliations:** Systems Biology of the Stress Response, Department of Biology, Technical University Darmstadt, Schnittspahnstraße 13, 64287 Darmstadt, Germany; Self-organizing Systems, Department of Electrical Engineering and Information Technology, Technical University Darmstadt, Merckstrasse 25, 64283 Darmstadt, Germany; Centre for Synthetic Biology, Technical University Darmstadt, Germany

**Keywords:** Stochastic transcription, Bayesian inference, telegraph model, smFISH, p53 signalling

## Abstract

The tumor suppressor p53 plays a crucial role in maintaining genome integrity in response to exogenous or endogenous stresses. The dynamics of p53 activation are stimulus- and cell type-dependent and regulate cell fate. Acting as a transcription factor, p53 induces the expression of target genes involved in apoptosis, cell cycle arrest and DNA repair. However, transcription is not a deterministic process, but rather occurs in bursts of activity and promoters switch stochastically between ON and OFF states, resulting in substantial cell-to-cell variability. Here, we characterized how stimulus-dependent p53 dynamics are converted into specific gene regulation patterns by inducing diverse forms of DNA damage ranging from ionizing and UV radiation to clinically relevant chemotherapeutics. We employed single molecule fluorescence *in-situ* hybridization (smFISH) to quantify the activity of target gene promoters at the single-cell and single-molecule level. To analyse this comprehensive data set, we developed a new framework for determining parameters of stochastic gene expression by Bayesian inference. Using this combined theoretical and experimental approach, we revealed that features of promoter activity are differentially regulated depending on the target gene and the nature and extent of the DNA damage induced. Indeed, stimulus-specific stochastic gene expression is predominantly regulated by promoter activation and deactivation rates. Interestingly, we found that in many situations, transcriptional activity was uncoupled from the total amount of p53 and the fraction bound to DNA, highlighting that transcriptional regulation by p53 is a multi- dimensional process. Taken together, our study provides insights into p53-mediated transcriptional regulation as an example of a dynamic transcription factor that shapes the cellular response to DNA damage.

## Introduction

Throughout their lives, cells in our bodies are confronted with various sources of stress. Altering gene expression is crucial for them to adapt to these stresses and elicit appropriate cell fate decisions. These changes in expression are mediated by the activity of transcription factors that change over time in response to cellular signal processing, thereby enabling the transfer of information from an incoming input into a cellular response. A central hub of the cellular stress response is the tumor suppressor p53. As one of the most studied transcription factors, p53 plays a pivotal role in maintaining genome stability and inhibiting tumorigenesis. Under normal conditions, p53 is modified by the ubiquitin ligase Mdm2 and degraded through the proteasome. In response to diverse stress signal, p53 undergoes post-translational modifications that stabilize the protein (1). After translocation to the nucleus, p53 promotes the expression of target genes that mediate its biological responses and determine cell fate decisions (**Figure 1A**) (2). Among the target genes of p53 are MDM2 and other negative regulators such as the phosphatase PPM1D/Wip1. These interactions form feedback loops that shape the dynamics of p53. Time-resolved analysis in individual living cells demonstrated that upon induction of DNA double strand breaks (DSBs), p53 accumulates in pulses of uniform amplitude and duration that are generated by excitability in the p53 network and cell-specific activation thresholds (3,4). However, in response to other stresses, p53 shows different temporal patterns of accumulation, such as a single p53 pulse following UV radiation (5), or a monotonic increase correlated with cell death upon cisplatin treatment (6).

**Figure 1.**
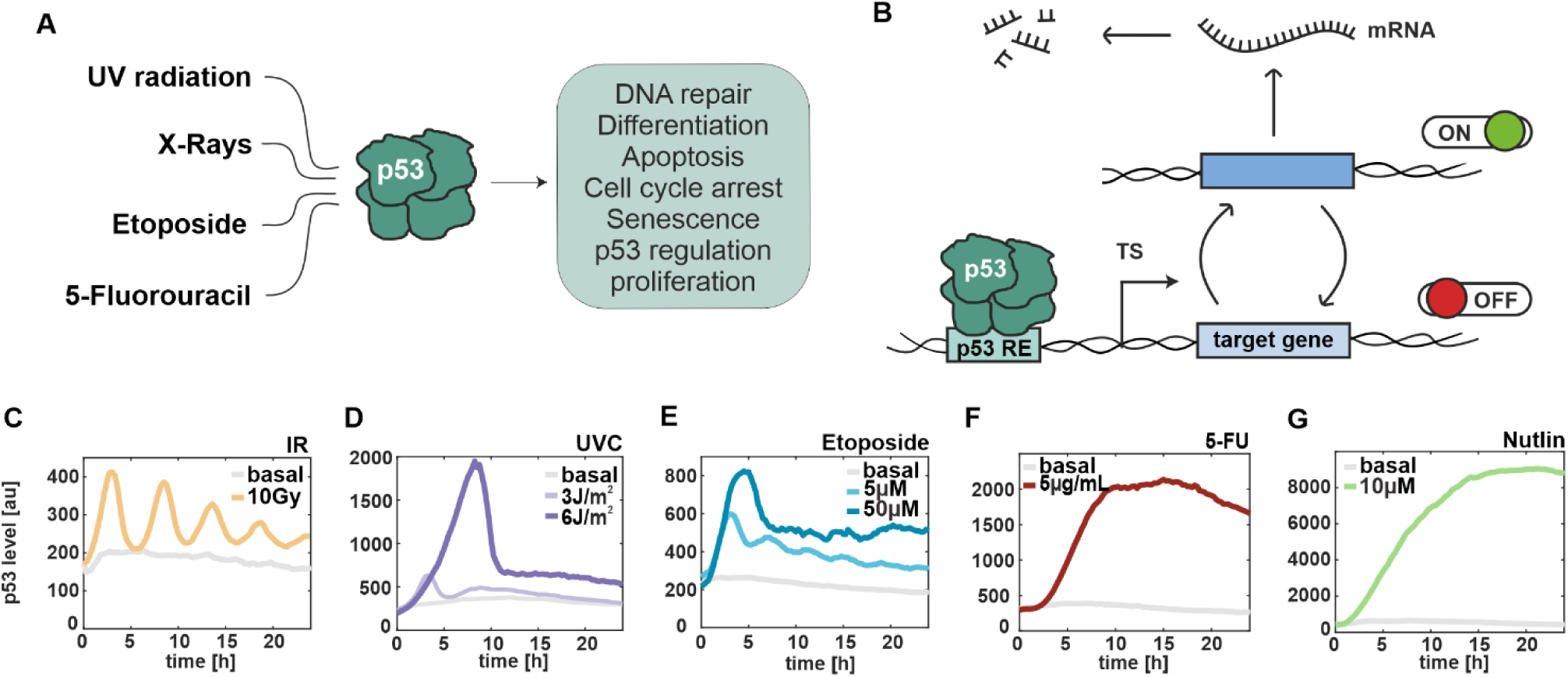
p53 dynamics are stimulus- and dose-dependent. (**A**) Schematical representation of selected stimuli, which induce the p53 stress response. (**B**) Illustration of promoter activity according to the random telegraph model. TS: transcription site, RE: response element (**C-G**) Median trajectories of p53 dynamics using live cell time-lapse microscopy of A549 reporter cell line. p53 levels are quantified as fluorescent intensity (y-axis) over 24 hours (x-axis). Basal levels are shown in grey. Cells were irradiated with ionizing radiation at 10 Gy (**C**) and UV radiation at 3J/m^2^ and 6J/m^2^ (**D**), and treated with different chemotherapeutics, such as 5µM and 50µM Etoposide (**E**), 5µg/mL 5-FU (**F**) and 10µM Nutlin-3a (**G**). More than 1000 cells were analysed for each condition.

When p53 induces expression of its target genes, a series of molecular events are triggered at the promoter. These include exchange of repressive to permissive histone marks, recruitment of the basal transcription machinery, modification of the C-terminal domain of RNA Polymerase II (Pol II) and elongation of nascent transcripts until a poly-A signal is reached at the end of the gene (7). Studies at the single molecule level have shown that transcription is a discontinuous process and occurs in episodic bursts of transcribed RNA (8), leading to cell-to-cell variability of RNA levels (9). To describe burst dynamics and their regulation, one of the most successfully applied models in both prokaryotes and eukaryotes is the random telegraph model (10). According to this model, a promoter switches stochastically between a transcriptionally active “ON” state, during which RNAs are produced by trains of Pol II, and an inactive “OFF” state, when no new RNA is transcribed, while existing mRNA undergoes degradation (**Figure 1B**). Within this framework, the telegraph model helps to elucidate diverse RNA distributions, enables rapid reliable parameter inference (11), and predicts rates at which genes are turned on and off by TFs, providing insights into gene expression regulation (12). Although more complex models, such as three-state and multi-state promoters, have been proposed, the telegraph model remains a robust framework for capturing the distributions observed in these more intricate systems (11). Parameter inference is typically performed using a Bayesian approach (13) and requires single-cell data at multiple time points. Consequently, single molecule fluorescence *in situ* hybridization (smFISH) (8,14,15), single-cell RNA sequencing (scRNA-seq) (16,17) and live-cell imaging using the MS2-MCP system (17, 18, 19) are among the most suitable experimental approaches for inferring the kinetic parameters of bursty transcription.

In our study, we implemented dual-color smFISH, utilizing two probe sets labelled with different fluorophores to simultaneously detect exons and introns of the same gene (8). This approach not only provided single-molecule resolution, but also enabled us to distinguish and quantify mature RNA in nucleus and cytoplasm as well as nascent RNAs at transcription sites (TSs). Unlike previous studies (8,21), we used this powerful tool in combination with a novel framework for describing stochastic gene expression based on Bayesian inference. We extended the set of considered measurable quantities representing statistical features of the telegraph model compared to the approach in Friedrich *et al*. (21) with the goal to improve parameter identifiability. We derived an explicit expression for the distribution of the RNA counts in polyploid cells and introduced an additional, phenomenological parameter to capture overdispersion in the fluorescence intensity from nascent RNA at single TSs. Bayesian inference estimates the distribution of model parameters given the measured quantities. This estimate is called the posterior distribution and provides a principled way to capture uncertainty in the model parameter (22). In higher-dimensional cases, where normalization of and thus direct sampling from the posterior distribution is not viable, Markov-chain-Monte-Carlo (MCMC) has become the baseline method of choice to obtain samples from the posterior distribution (23). We thus used MCMC to update our knowledge of model parameters in the presence of the experimentally acquired data.

Quantification of bursting transcription of p53 target genes has been previously explored. In 2019, Friedrich *et. al* used an smFISH-based approach and demonstrated the p53-dependent transcription is intrinsically stochastic and mainly regulated by the fraction of active promoters (burst frequency) (21). Similarly, Hafner *et. al* observed in living cells that p53 predominantly influences the probability of transcriptional activation, likely due to transcriptional saturation during the ON state (24). Both studies reported modulation of transcriptional dynamics following DNA damage induced by ionizing radiations (IR). However, the regulation of p53 target gene promoters in response to other external stimuli remains unclear.

Our study aimed to systematically compare p53-dependent gene expression in a subset of targets upon various clinically relevant treatments and to investigate transcriptional regulation at the single-molecule and single-cell levels. We provide evidence for an uncoupling between p53 dynamics and its target gene expression. Additionally, using a combinatorial experimental and mathematical approach we identified that regulation of promoter activity is both stimulus- and gene-specific, and mainly driven by the frequency of promoters switching between ON and OFF states.

## Materials and Methods

### Cell line

A549 cells (#CCL-185) were cultured in McCoy’s 5A growth medium (GE Healthcare, Solingen, GER) supplemented with 10% (v/v) Fetal Bovine Serum, 2mM GlutaMAX, 100 U/mL penicillin and 100 µg/mL streptomycin. When required, the medium was supplemented with selective antibiotics to maintain transgene expression (400 μg/ml G418, 50 μg/ml hygromycin, or 0.5 μg/ml puromycin). The A549 p53 reporter and p53 knockdown cell lines have been described before by Finzel *et al,* 2016 (25).

### Irradiation and treatments with chemotherapeutics

Cells were irradiated with 10 Gy X-rays with a dose rate of 1 Gy/26 s and an energy of 250 keV. UV-C irradiation (256 nm) was performed using a UV lamp at a 40 cm distance, with exposures of 2 s (3 J/m²) and 4 s (6 J/m²). Prior to UV-C irradiation, the cell culture medium was replaced with sterile PBS. Consequently, smFISH measurements distinguished between basal conditions (no medium change) and the 0 h time point (PBS replacement). Drug treatments were applied at the following final concentrations: etoposide (#E1383, Sigma-Aldrich, St. Louis, USA) at 5 µM (low) and 50 µM (high), 5-fluorouracil (CAS 51-21-8, Sigma-Aldrich, St. Louis, USA) at 5 µg/mL, and Nutlin-3a (#N6287, Sigma-Aldrich, St. Louis, USA) at 10 µM.

### Live-cell time lapse microscopy

A549 reporter cells with both p53-mVenus and H2B-CFP nuclear marker were used for live-cell time-lapse microscopy. Two days prior the experiment, cells were grown on 24-well clear bottom plates or 35-mm clear bottom plates (both Ibidi GmbH, Gräfelfing, GER) depending on the treatment. Medium was changed to FluoroBrite^TM^ DMEM microscopy medium (Thermo Fisher Scientific, Waltham, MA, USA) supplemented with 5% Fetal Bovine Serum and cells were kept at stable conditions of 5% CO2 and 37°C. Cells were imaged in a Nikon Eclipse Ti-E inverted microscope with perfect focus system (Nikon) using a 20× CFI Plan Fluor objective (NA 0.75) and a Nikon DS-Qi2 digital camera. X-Cite 120 LED illumination system (Lumen Dynamics) was used for imaging and appropriate filter sets: (mVenus: 500/20 nm excitation (EX), 515 nm dichroic beam splitter (BS), 535/30 nm emission (EM); eCFP: 436/20 nm EX, 455 nm BS, 480/40 nm EM. Images were acquired every 15 minutes for 24h.

### Automated tracking of cells and analysis of p53 dynamics

Automated segmentation of nuclei and quantitative analysis of p53 levels were performed in MATLAB (MathWorks, Natick, USA) using custom written scripts as described in Finzel *et al*. 2016 (25). In brief, flat field correction and background subtraction were applied to raw images, afterwards individual nuclei were segmented using thresholding and watershedding algorithms. Segmented cells were then assigned to corresponding cells in following images using a greedy match algorithm. Only cells tracked from the first to last time-point were considered for quantitative analysis. The integrated nuclear fluorescence intensity was used to determine p53 abundance and generate single-cell trajectories for individual cells over time.

### Single molecule FISH

Stellaris probe sets for single-molecule fluorescent in situ hybridization (smFISH) (Biosearch Technologies) were custom-designed for targeting p53 target genes and were directly adopted from Friedrich *et al.* 2019 (21). Exon probes were labelled with CAL Fluor Red 610, while intron probes with Quasar 670 dye. A549 wild-type cells were grown for two days on 18mm uncoated coverslips (thickness #1). After treatment, cells were washed with sterile ice-cold PBS at indicated time points, fixed with 2% paraformaldehyde (EM-grade) (Electron Microscopy Sciences, Hatfield, USA) for 10 min at room temperature and permeabilized over night with 70% ethanol at 4 °C. Each probe set was pre-diluted in Tris-EDTA (Sigma Aldrich, St. Louis, USA) and hybridized at a final concentration of 0.1 μM for 16h at 37°C in a humified chamber. Afterwards, cells were washed and incubated with Hoechst-33342 (Life technologies, Carlsbad, USA) for 10 min at room temperature for nuclear staining and with HCS CellMask™ (Thermo Fisher Scientific, Waltham, MA, USA) for cytoplasmic counterstain. Coverslips were then mounted on Prolong Gold Antifade (Life Technologies). Cells were imaged on a Nikon Eclipse Ti-E inverted fluorescence microscope with a DS-Qi2 camera. A 60x plan apo objective (NA 1.4) and appropriate filter sets were used: (Hoechst: 387/11-nm excitation (EX), 409-nm dichroic beam splitter (BS), 447/60-nm emission (EM); Alexa Fluor 488: 470/40 nm (EX), 495 nm (BS), 525/50 nm (EM); CAL Fluor Red 610: 580/25 nm (EM), 600 nm (BS), 625 nm (EX); Quasar 670: 640/30 nm (EX), 660 nm (BS), 690/50 nm (EM)). Images were acquired as multipoints of 21 z-stacks with 300 nm step-width using Nikon Elements software (Nikon instruments, Tokyo, JPN).

### Analysis of smFISH data in *FISH-Quant*

Automated segmentation of nuclei and cytoplasm was performed using Cellpose (26). TSs were manually detected based on the intron signal in nuclei in all z-planes. In brief, according to the *FISH-Quant* workflow as described in Mueller *et al,* 2013 (27) for spot detection, images were filtered, pre-detection was performed, then spots were fitted, and fits were further thresholded to exclude outliers. TS detection was based on the intron signal and an average cytoplasmic spot intensity was computed for determining the number of nascent RNAs at the detected TS. Each analysis was performed using the *FISH-Quant* batch processing toolbox. RNA counts and detected TSs were then used for mathematical modelling.

### The Likelihood factors from Bayesian inference

The factors of the likelihood, abbreviated as *L1* and *L*2, are detailed in the following. The first factor *L1* accounted for the measured RNA count *m* (estimated from exon fluorescence) and the amount *k* < *K* of active TSs in a cell (with a total of *K* TSs in this cell, active or inactive). We assumed the decomposition *L*_1_ = *L*_1,1_ ⋅ … ⋅ *L*_1,*C*_, where the cells (enumerated 1, …, *C*) were considered independent. Based on the telegraph model, *L*_1,*c*_ then determines the likelihood of seeing a total of *m* RNA molecules at steady-state when *K* independent telegraph models with identical parameters *θ* = (*λ*, *γ*, *μ*, *δ*) are present in cell *c*, of which *k* are active and *K* − *k* are inactive. Let *X*_*n*_ ∈ {0, 1} give the activity of the *n*-th TS and *Y*_*n*_ ∈ {0, 1, 2, … } give the amount of present RNA from the *n*-th TS. Then, with *X* = *X*_1_ + ⋯ + *X*_*K*_ and *Y* = *Y*_1_ + ⋯ + *Y*_*K*_, the likelihood *L*_1,*c*_ was obtained in closed form as

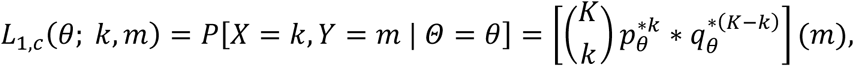

where the superscript *x*^∗*k*^ denotes the *k*-times convolution of a function *x* with itself and the partial mass functions *p*_*θ*_ = *P*[*X*_*n*_ = 1, *Y*_*n*_ = *m*] and *q*_*θ*_ = *P*[*X*_*n*_ = 0, *Y*_*n*_ = *m*] (equal for all *m*) were found to be given as

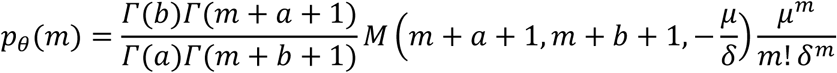

and

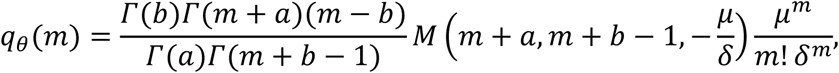

where *M* denotes Kummer’s confluent hypergeometric function and *a* = *λ*⁄*δ* and *b* = (*λ* + *γ*)⁄*δ*. The derivation of the equation for *L*_1,*c*_ can be found in the Supplementary Methods and is based on a result by Peccoud *et al.* (*28*). In this likelihood alone, the parameters stay ambiguous w.r.t. their scale, because of the consistent division by *δ*.

The second factor *L*2 accounted for the measured fluorescence from exon probes in the vicinity of a detected TS, but only in those cases in which fluorescence from intron probes was present as well. Presence of fluorescence from intron probes was used to declare the respective TS as active. We assumed the decomposition *L*_2_ = *L*_2,1_ ⋅ … ⋅ *L*_2,*T*_, where the active TSs (enumerated 1, …, *T*) were considered independent. The factor *L*2,t then determines the likelihood that the measured total exon fluorescence *z* for the detected active TS *t* is a sample from a Gamma distribution, i.e.

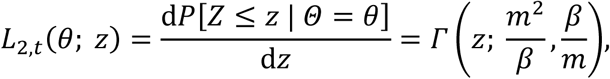

where *Γ* is the Gamma probability density function in shape-scale parametrization, obtained by moment-matching with the statistics *m* (mean) and *β* (variance). The mean *m* of the distribution was chosen as the total exon fluorescence intensity from nascent RNA currently transcribed at an active TS, as predicted by the telegraph model with parameters *γ*, *λ* and *μ*. This mean was obtained by rejection sampling from the telegraph model in the following way. We first sampled *N* trajectories *π*_1_, …, *π*_*N*_ from the underlying telegraph process determining active and inactive periods of the TS.

Conditioned on a trajectory *π*_*n*_, we sampled an inhomogeneous Poisson point process (with arrival rates in {0, *μ*}, depending on the active and inactive periods in *π*_*n*_) to obtain a sample of positions *p*_*n*_ = (*p*_*n*,1_, *p*_*n*,2_, …) of polymerases on the gene. For a sample of polymerase positions *p*_*n*_, we then calculated the exon fluorescence intensity *z*_*n*_ of nascent RNA if at least one polymerase resided within the range of an intron of the respective gene downstream of at least one corresponding intron probe or otherwise rejected the sample (*π*_*n*_, *p*_*n*_) and resampled it. The elongation rate of the polymerases was considered a hyperparameter and set to 50bps, the transcription initiation rate was identified with the burst rate μ. The mean *m* was then approximated as

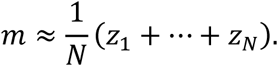

The variance *β* of the Gamma distribution was set as an additional model parameter that is not mechanistically justified but considered a phenomenological parameter.

### Configuration of the MCMC

The initial values of the model parameters were manually adjusted to reasonable values. Since all parameters were positive, real, and could be of different orders of magnitude, inference was done on the logarithmic values and the logarithmic step proposal distribution was chosen to be a multivariate normal distribution with initial unit covariance (corresponding to a multivariate log-normal scaling- distribution in the linear domain). A Robbins-Monro schedule was used to adjust the covariance continuously during the burn-in period from a moving window of samples (29). The schedule and the length of the moving window were considered hyperparameters and adjusted manually to achieve allowable acceptance rates during the posterior sampling phase.

### Chromatin Immunoprecipitation

A549 wild-type cells were grown in 10-cm plates to reach 80% confluency. After treatment, cells were washed with sterile ice-cold PBS at indicated time points, fixed with 1% paraformaldehyde (EM-grade) (Electron Microscopy Sciences, Hatfield, USA) for 10 min at room temperature and incubate with 125 mM Glycine in PBS for 5 min to stop fixation. Cells were washed with ice-cold PBS and harvested in PBS supplemented with 1 mM PMSF. Cell pellet was resuspended in Lysis buffer (5 mM Tris–HCl, pH 8.0, 85 mM KCl, 0.5% Igepal-CA630 supplemented with protease inhibitor cocktail and 1 mM PMSF) and incubated for 20 min on ice. Nuclear pellet was collected by centrifugation, resuspended in Sonication buffer (50 mM Tris–HCl, pH 8.1, 0.3% SDS [w/v], 10 mM EDTA supplemented with 1 mM PMSF and Protease Inhibitor Cocktail), and incubated for at least 30 min on ice. Chromatin fragmentation was performed using the Bioruptor® Pico (Diagenode,LLC, NJ, USA) according to the manual. Two cycle of sonication (30 sec ON, 30 sec OFF) per 6 × 10^6^ cells were sufficient to ensure fragment size of 200-700 bp. 10 µg of chromatin was then diluted with Dilution buffer (16.7 mM Tris–HCl, 167 mM NaCl, 0.01% SDS [w/v], 1.2 mM EDTA, 1.1% Triton [v/v], 1 mM PMSF, Protease Inhibitor Cocktail) and incubated overnight at 4°C with 1 µg of p53 antibody (#2527, Cell Signaling Technology, MA, USA) or a control IgG (Normal rabbit IgG, Merck Millipore). To collect the immunocomplexes, 20 μl of Dynabeads Protein G (Thermo Fisher Scientific) were added for 2 h at 4 °C. The beads were washed once with low salt washing buffer (0.1% SDS [w/v], 2 mM EDTA, 20 mM Tris–HCl pH 8.1, 1% Triton X-100 [v/v], and 150 mM NaCl), high salt washing buffer (0.1% SDS [w/v], 2 mM EDTA, 20 mM Tris–HCl pH 8.1, 1% Triton X-100 [v/v], and 500 mM NaCl) and LiCl washing buffer (10 mM Tris–HCl pH 8.1, 1 mM EDTA, 1% IGEPAL CA630 [v/v], 1% deoxycholic acid [w/v], 250 mM LiCl), and TE-Buffer (10 mM Tris– HCl pH 8.1, 1 mM EDTA). Chromatin was eluted from the beads for 30 min at 65°C in Elution buffer (1% SDS, 0.1 M NaHCO3) twice. Crosslinks were reversed incubating the samples with 200 mM NaCl at 65°C overnight and by following addition of 10 µg/mL RNase A for 30 min at 37°C and 100 μg/ml Proteinase K, 10 mM EDTA, and 40 mM Tris–HCl pH 6.5 for 3 h at 45°C. Finally, the DNA was purified using the Monarch® PCR & DNA Cleanup Kit (NEB, Ipswich, MA) and 3 μl of each sample was used for qPCR.

### Quantitative Real-time PCR (RT-qPCR)

RNA was extracted at the indicated time points using the Monarch® Total RNA Miniprep Kit (#T2010, NEB). cDNA was generated from 1 µg extracted RNA using M-MuLV reverse transcriptase and oligo-dT primers (both NEB). Quantitative real-time PCR was performed in triplicates using ProtoScript II Reverse Transcriptase (NEB) and SYBR Green reagent (Thermo Fisher Scientific, USA) on a StepOnePlus PCR machine (Thermo Fisher Scientific, USA). Relative gene expression changes were analyzed using the 2^−ΔΔCT^ method. Actin served as the housekeeping gene. The final concentration of used primers was 243.2 nM. The following intron-spanning primers were used: BAX forward: CTGACGGCAACTTCAACTGG; BAX reverse: GATCAGTTCCGGCACCTTGG; MDM2 forward: AGATGTTGGGCCCTTCGTGAGAA; MDM2 reverse: GCCCTCTTCAGCTTGTGTTGAGTT; CDKN1A forward: TGGACCTGTCACTGTCTTGT; CDKN1A reverse: TGGACCTGTCACTGTCTTGT; PPM1D forward: ATAAGCCAGAACTTCCCAAGG; PPM1D reverse: TGGTCAATAACTGTGCTCCTTC; β-ACTIN forward: GGCACCCAGCACAATGAAGATCAA; β-ACTIN reverse: TAGAAGCATTTGCGGTGGACGATG. For ChIP assays, these primers were used: BAX forward: AACCAGGGGATCTCGGAAG; BAX reverse: AGTGCCAGAGGCAGGAAGT; MDM2 forward: GTTCAGTGGGCAGGTTGACT; MDM2 reverse: CGGAACGTGTCTGAACTTGA; CDKN1A forward: AGCCTTCCTCACATCCTCCT; CDKN1A reverse: GGAATGGTGAAAGGTGGAAA.

### Western blot

Cells were grown for 2 days in 6 cm plates to reach 80% density. After treatment, cells were harvested at indicated time points. Protein isolation was performed by lysis in RIPA buffer in the presence of protease and phosphatase inhibitors (Carl Roth GmBH, Karlsruhe, GER and Sigma Aldrich, St. Louis, USA) as well as Panobinostat (#404950-80-7, Cayman Chemical, USA) as deacetylase inhibitor. For the SDS-PAGE, 20 µg of protein were used and loaded on 10% acrylamide gels. Afterwards, proteins were transferred to PVDF membranes (GE Healthcare) by electroblotting (Bio-Rad). The membrane was blocked with 5% milk in TBS-T and incubated overnight with the primary antibody. The next day, the membrane was washed with TBS-T, incubated with secondary antibody coupled to peroxidase, washed again, and protein levels were determined using chemoluminescence (WesternBright® Quantum®, Advansta). For detection of p53 acetylation, all blocking, wash and incubation buffers contained Panobinostat. Precision Plus Protein Dual Color Standard (Bio-Rad, Feldkirchen, GER) was used for molecular mass comparison and GAPDH as loading control. The following antibodies were used: anti-GAPDH (Sigma-Aldrich, G9545), anti-p53 (Santa Cruz #DO1, Cell Signaling, #9282), anti-p53K382ac (Abcam, ab75754), goat anti-rabbit-IgG-HRP (Thermo Fisher Scientific), goat anti-mouse-IgG-HRP (Thermo Fisher Scientific).

### Labelling of cells with EdU

A549 wild type cells were seeded on 15mm circular, uncoated, #1 high-precision coverslips (round) two days prior the experiment. After treatment, they were incubated with 1mM EdU for 1h and subsequently washed with PBS. Cells were then fixed with 2% paraformaldehyde for 10 min at room temperature and permeabilized with 0.1% Triton X-100 in PBS for 20 minutes. After washing, cells were stained with 20 µM Eterneon Red 645-Azide according to the manual using the EdU Click-647 ROTI®kit for Imaging (Carl Roth, #7777.1). Counterstain of nuclei was performed using Hoechst- 33342. Coverslips were finally mounted and imaged as described for the smFISH method.

## Results

### 1. P53 dynamics are stimulus- and dose-dependent

To investigate stimulus-dependent gene regulation, we induced different types of DNA damage, focusing on ionizing and UV radiations as well as clinically relevant chemotherapeutics. First, we characterized p53 dynamics in response to different doses of these stimuli in a previous established small cell lung carcinoma cell line A549 expressing wild type p53 fused with the mVenus fluorescent protein (30) using live-cell time-lapse microscopy (**Figure 1C-G**). Cells exposed to ionizing radiation (IR) accumulate DSBs, triggering the DNA damage response (DDR) primarily mediated by the PI3K- like kinase ATM (31). A subsequent phosphorylation cascade induces p53 accumulation in the nucleus in pulses of uniform amplitude and duration (32). We observed these pulsatile p53 dynamics after irradiation with 10 Gy (**Figure 1C**). In contrast, UVC light provokes cross-link of consecutive pyrimidine bases leading to exposure of single-stranded DNA (ssDNA) and activation of an ATR-dependent DNA damage response (33). Upon low doses of UV, p53 exhibited pulsatile dynamics similarly to the response upon IR. Upon high doses, p53 showed instead a single pulse of accumulation with higher amplitude and duration (**Figure 1D**). A bimodal response was also observed upon etoposide treatment. Indeed, at 5 µM concentration, oscillatory p53 dynamics were detected, while at higher concentrations p53 accumulated in a single pulse (**Figure 1E**). This chemotherapeutic drug stabilizes the binding between DNA topoisomerase II and DNA resulting in DSBs (30). In contrast, the antimetabolite 5-fluorouracil (5-FU) mainly acts by blocking the enzyme thymidylate synthase and by inhibiting both RNA and DNA synthesis (34). After treatment with 5- FU, p53 levels constantly increased over time. Decreasing p53 at the end of the experiment might be due to cytotoxicity (**Figure 1F**). To complete our analysis, we induced p53 accumulation in absence of DNA damage using Nutlin-3a as an inhibitor of the p53-Mdm2 interaction (35). As expected, we observed a constant increase of p53 levels over time (**Figure 1G**). Moreover, we noted substantial differences in p53 accumulation across treatments, with the highest levels detected following Nutlin, 5-FU, and high-dose UV radiation. We validated these stimulus-specific p53 dynamics using Western Blot assays in wild-type A549 cells, which were used to quantify transcription in subsequent experiments (**Supplementary Figure 1).** In conclusion, we identified a set of stimuli that resulted in different dose-dependent dynamics of p53 in A549 cells that allowed us to investigate the correlation between temporal changes in transcription factor levels and stochastic expression of its target genes.

### 2. Expression of p53 targets show stimulus and gene-specific pattern

In a previous study, we demonstrated that following IR, the first p53 pulse induces a general increase in target gene expression (21). However, at later time points, gene-specific changes were observed. If p53 exhibits different dynamic patterns, do target genes adjust their expression accordingly? And if so, how do these changes manifest over time? To address these questions, we selected four well-characterized p53 target genes involved in different cell fate programs: MDM2, PPM1D/ Wip1 (both feedback regulators); CDKN1A/ p21 (cyclin-dependent kinase inhibitor) and BAX (apoptosis). We first confirmed that expression of the chosen target genes was induced by p53 accumulation in response to the selected stimuli by comparing mRNA levels in populations of p53 wild type and knockdown cell lines using quantitative RT-qPCR (**Supplementary Figure 2A**). We then employed smFISH (8,21) and measured total RNA levels over time using exon-targeting probes (**Figure 2A- B**). Nuclear and cytoplasmic stainings (**Supplementary Figure 2B**) allowed us to identify individual cells and assign mRNAs to both subcellular compartments using FISH-Quant (27) and custom-written Matlab scripts (see Methods for details). smFISH requires cell fixation at defined time points, which were selected based on changes of p53 levels in response to different doses of the previously described stimuli (**Figure 2C).**

**Figure 2.**
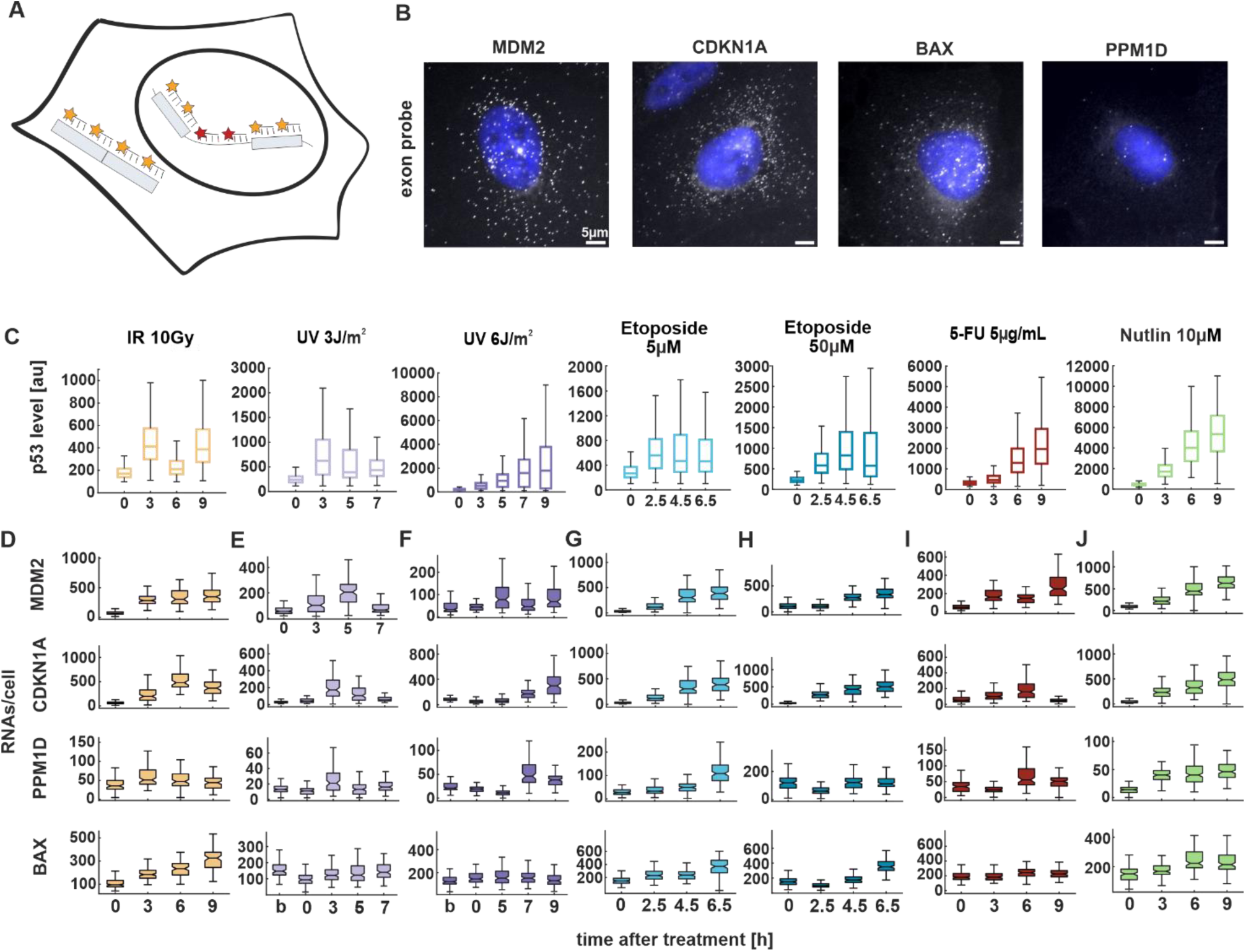
p53 target genes show stimulus and gene-specific expression patterns. (**A**) Schematic depiction of smFISH probe sets. Exon (yellow) and intron (red) probes were labelled with the fluorophores CAL Fluor Red 610 and Quasar 670, respectively. (**B**) Fluorescence microscopy images of smFISH probes CAL Fluor 610 (grey) overlayed with Hoechst 33342 staining (blue) 3 h after Nutlin treatment for the indicated target genes in A549 cells. Scale bar corresponds to 5 μm distance; images were contrast and brightness enhanced for better visualization. (**C**) Median p53 levels from live-cell time-lapse microscopy experiments. Boxplots represent the quantification of nuclear fluorescent intensity at indicated time points following treatments (**D**) RNAs per cell were quantified using FISH-Quant for four target genes at the indicated time points upon treatment with the selected stimuli and displayed as box plots. For all boxplots, lines indicate medians of distributions; boxes include data between the 25^th^ and 75^th^ percentiles; whiskers extend to maximum values within 1.5× the interquartile range. Approximately 100 cells per experiment were analyzed (see Supplementary Table 1 for details).

Surprisingly, our smFISH analysis revealed a disconnect between p53 dynamics and mRNA expression (compare **Figure 2C and D-J**). Specifically, following irradiation with 10 Gy, we observed pulsatile p53 dynamics which did not correspond to the expression patterns of its target genes. The selected genes exhibited either transient expression or a sustained increase in RNA levels over time (**Figure 2D**). In contrast, after UV radiation at 3J/m^2^, only a single full p53 pulse was detected within the analyzed time points. MDM2, CDKN1A, and BAX initially showed increased RNA levels which returned to basal levels at later time points (**Figure 2E**). At higher UV doses, although p53 levels progressively increased, RNA levels rose only at very late time points (**Figure 2F**). Notably, BAX showed no induction of gene expression at both low and high UV doses (**Figure 2E-F**). Following treatment with etoposide, p53 levels were twice as high when cells were treated with 50 μM compared to 5µM. However, the expression pattern of all target genes remained unchanged between the two conditions, with RNA levels steadily increasing over time (**Figure 2G- H**). Upon 5-FU treatment, p53 accumulation increases monotonously. Surprisingly, target gene expression did not reflect these dynamics, as we observed only slight changes in RNA levels for all target genes (**Figure 2I**). As expected, target genes showed increased expression following Nutlin treatment (**Figure 2J**). Interestingly, we noted that basal RNA levels varied among the genes, with BAX showing high number of transcripts already without induction. Moreover, each p53 target reached its maximum level of transcription at different time points after induction, defining a gene-specific pattern of expression depending on the stimulus (**Supplementary Figure 3**). Overall, our findings indicate that RNA expression is gene-specific upon most of the stimuli, but poorly dose-dependent. Moreover, transcription patterns do not correlate with p53 dynamics over the selected time points.

Surprisingly, most target genes do not exhibit a significant increase in expression following UV radiation, especially at high doses, despite elevated p53 levels. Several studies have reported that UV exposure leads to transcriptional inhibition through both elongation blockage and initiation impairment (36). To assess this phenomenon, we imaged EU-labeled RNA levels in cells at 3h post-irradiation. Consistent with expectations, our analysis revealed a reduction in nascent RNA levels upon both low and high doses of UV compared to unirradiated cells (**Supplementary Figure 4**).

### 3. Identification of regulation patterns of promoter activity through Bayesian inference

While RNA transcript measurements provide valuable insights into gene expression, they are insufficient to fully characterize the bursting dynamics of target gene promoters. Therefore, we employed dual-color smFISH labeling to measure both mature RNAs (as described above) and nascent transcripts using intron-targeting probes conjugated to a distinct fluorophore (**Supplementary Figure 2B**). This approach enabled the identification of actively transcribing transcription sites (TSs), which, as expected, were detected exclusively in the nucleus. Because our study was conducted in a polyploid cell line, we identified more than two TSs for all the selected p53 targets. The maximum number of genomic loci has been previously validated by Friedrich *et al.*(21). Furthermore, our data confirm independent allele activity, as we observed single cells displaying zero, one, two, or more active TSs (**Supplementary Figure 5A**). Measurements of RNA molecules, TS counts and fluorescence intensities of the exon probes were extracted from the FISH-Quant analysis (27) (see Methods for details). When quantifying nascent RNAs, it is important to keep in mind that individual oligonucleotides of the exon and intron probe sets are unevenly distributed across the gene length (**Supplementary Figure 5B**). Therefore, the fluorescence intensity of individual nascent transcripts depends on the respective progression of RNA polymerase II (Pol II).

To derive bursting parameters, mathematical modelling based on mRNA distributions and polymerase occupancies has been previously used (8,21). However, this approach relies on Poisson-like distributions and provides limited details in the reported parameters. To achieve more accurate estimates of transcriptional activity for p53 target gene promoters, we refined a framework based on Bayesian inference and the random telegraph model. This model consists of a non-leaky two-state promoter, switching between active and inactive periods with activation rate *λ* and deactivation rate *γ*. During an active period, RNA is transcribed at the promoter with a rate *μ*. Existing RNA is degraded with rate *δ* (**Figure 3A**). Investigation of the statistical relation between the telegraph model and nascent RNA fluorescence revealed an overdispersion of the fluorescence intensity data, which was captured by an additional model parameter *β*.

**Figure 3.**
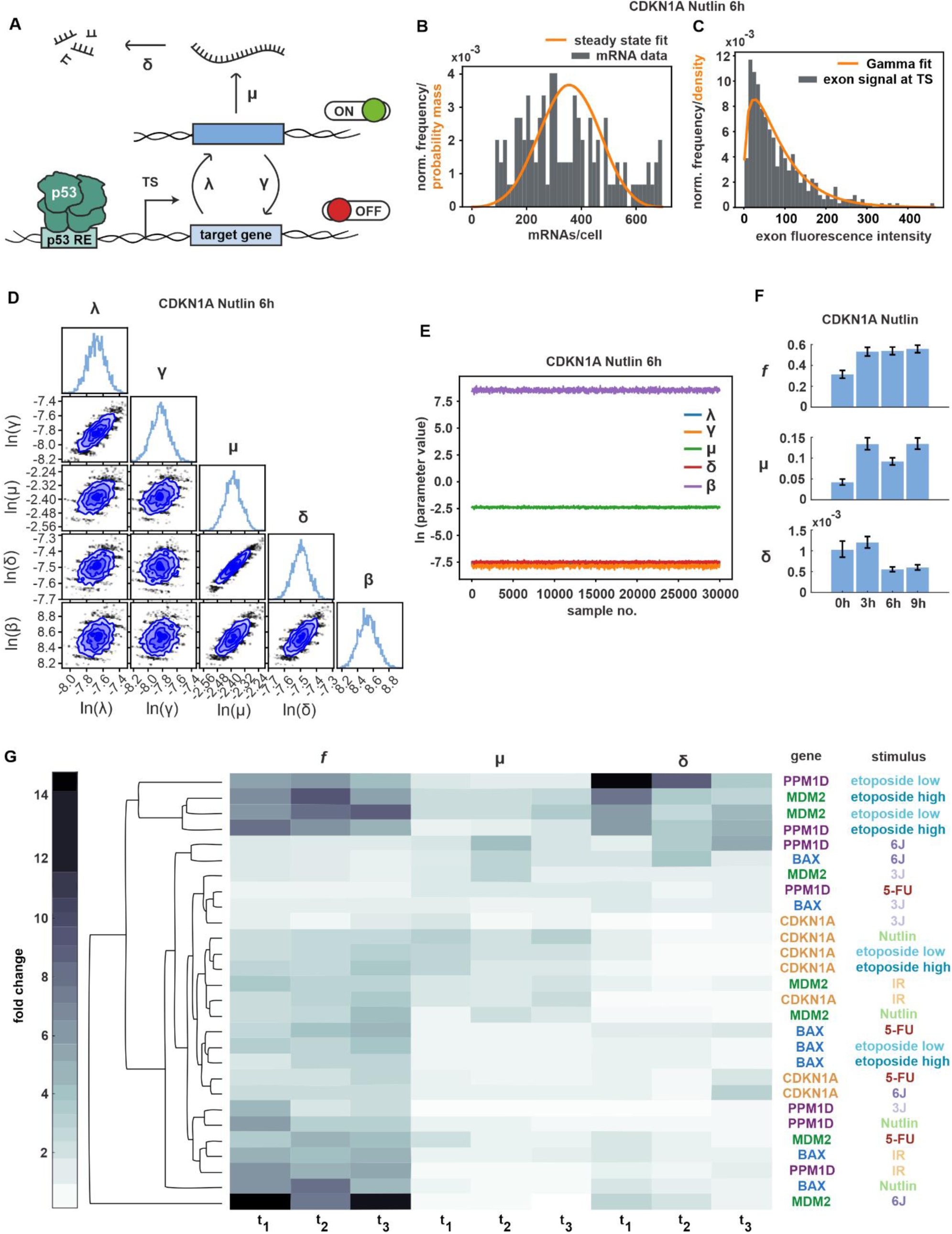
Bayesian inference enables characterization of promoter features. (**A**) Schematic representation of promoter activity based on the random telegraph model, incorporating RNA production and degradation. Transcriptional parameters were inferred using Markov-Chain-Monte-Carlo (MCMC): activation rate λ, deactivation rate γ, transcription rate µ, degradation rate δ, variance of nascent RNA fluorescence intensity *β* (not shown). (**B**) Histogram of CDKN1A mRNA transcripts at 6h time point upon Nutlin treatment (grey bars). The orange curve represents the predicted probability mass function by the model for multiple independent TSs. (**C**) Histogram of measured fluorescence intensity levels at TS for CDKN1A at 6h time point following Nutlin treatment (grey bars). The orange curve represents the density of the fitted Gamma distribution. (**D**) Corner plot displaying low-dimensional marginals of the MCMC-obtained empirical estimate of the posterior distribution in the (natural) logarithmic domain. The diagonal elements show the marginal distributions of individual parameters, while the off-diagonal elements present two-dimensional marginal distributions for parameter pairs. (**E**) Plot of posterior sample traces for all parameters from the MCMC run for CDKN1A at 6h following Nutlin treatment. Convergence of the Markov chain was assessed visually. λ values (in blue) are not visible because they overlap with γ and δ values. (**F**) Bar plot depicting the posterior mean estimates of *f* (corresponding to λ/(λ+γ)), µ, and δ for a representative example. The horizontal axis represents time points after treatment, while the vertical axis shows posterior means and confidence intervals (2.5% and 97.5% quantiles). (**G**) Summary of MCMC estimates of *f*, µ, and δ represented as a clustergram. Each row corresponds to a specific gene and stimulus, while each column represents a given time point post-treatment (3h, 6h and 9h for IR, 5-FU and Nutlin; 2.5h, 4.5h and 6.5h for etoposide low and high; 3h, 5h, 7h, for UV 3J/m^2^ and 5h, 7h, 9h for UV 6J/m^2^). In the heatmap, color intensity represents fold changes, ranging from 0-fold (white) to 15-fold (black), normalized to the basal (untreated) condition. The dendrogram was generated by computing pairwise Euclidean distances between time points and applying the average linkage method.

The Bayesian posterior distribution is proportional to the product of the observation likelihood and the prior distribution over the model parameters, which in our case were *γ*, *λ*, *μ*, *δ* and *β*. Inference was done for each condition, gene, and time-point independently. An improper uniform prior over the non-negative real parameter space was chosen. The observation likelihood consisted of a product of two independent factors (given the model parameters, i.e. a sample thereof), of which more details can be found in the Methods section.

One of the factors accounted for was the total measured RNA counts (estimated from exon probes) and the total number of TSs present in a cell. Based on the telegraph model, we calculated the likelihood to measure the given total amount of RNA, when multiple independent telegraph models with identical parameters (*γ*, *λ* and *μ*, all normalized by *δ*) are present (**Figure 3B**).

The other factor accounted for was the measured fluorescence from exon probes in the vicinity of a detected TS, but only in those cases in which fluorescence from intron probes was present as well. Presence of fluorescence from intron probes was used to declare the respective TS as active. Having assumed a separation of timescales between spatial displacement and transcription of RNA, we declared the measured exon fluorescence to come from nascent RNA. We calculated the likelihood that the measured exon fluorescence intensities for the detected active TSs are samples of a Gamma distribution with the following statistics. Its mean was predicted by the telegraph model with parameters *γ*, *λ* and *μ,* and its variance was given by the additional phenomenological model parameter *β*. The positions of polymerases on the gene determine the count of exon probes already transcribed in the nascent RNAs that contribute to the fluorescence. These positions were inferred using the telegraph model. As each single exon probe contributes to the measured total intensity by a fixed amount, we could determine the mean fluorescence intensity at the active TSs from the mean occupancy of the gene by Pol II (**Figure 3C**).

Correlation in the posterior of the parameters varied across cases (conditions, genes, time-points). However, in most cases the strongest correlations were observed between *λ* and *γ* as well as *μ* and δ (**Figure 3D**). Hyperparameters were carefully calibrated for (single-chain) MCMC to appropriately sample the Bayesian posterior (details in the Methods section). All samples were obtained in the logarithmic domain. We generated 30,000 posterior samples with an additional 10,000 samples for the burn-in period. Convergence was later assessed visually from the sample traces (**Figure 3E**). All reported estimated parameters correspond to posterior mean estimates.

Using this combined experimental and theoretical approach, we generated a comprehensive data set describing gene- and stimulus-specific regulation of promoter activity by p53. For each of the stimuli described above, we selected informative time points and analysed hundreds of individual cells. For simplicity, we defined *f* as the fraction λ/(λ+γ) of active promoters in the steady state. Characterizing *f*, μ and δ provided insights into regulation of transcriptional bursting for p53 target genes. However, since δ is a derived parameter, its estimation should be interpreted with caution, as it may not directly reflect underlying biological processes.

We first analyzed stochastic gene expression from the CDKN1A promoter following Nutlin treatment. We observed that in this condition, *f* exhibited a two-fold increase at 3h post-treatment and remained constant over time. We observed a similar pattern for *μ*. In contrast, *δ* decreased at later time points (**Figure 3F**). To allow a better comparison of promoter regulation over time, we normalized each parameter to the basal time point and present the resulting fold-changes as a heat map. Using this approach, we analysed promoter activity of the CDKN1A gene upon the selected stimuli. We again observed that *f* was strongly regulated in most conditions, with additional contributions by *μ* (**Supplementary Figure 5C**). In this context, we determined the reproducibility of MCMC estimates on biological replicates. While the main findings were robust, we observed a certain degree of variability for some parameters and time points due to technical limitations as well as the stochastic nature of the biological process.

To systematically analyse promoter activity across genes and conditions, we performed hierarchical clustering of normalized transcriptional parameters (**Figure 3G**). This approach allowed us to gain insights into distinct transcriptional regulatory patterns. The underlying raw and processed measurements are available via our institutional repository (see data availability section). Our analysis revealed that p53 target gene expression is predominantly regulated by the switching ON/OFF (*f*) of their promoters, with a minor modulation of μ and δ. However, there appear to be both stimulus- and gene-specific features of promoter regulation. In absence of DNA damage, upon Nutlin treatment where transcriptional activation is driven solely by the accumulation of p53 over time, promoter properties became most easily apparent. Specifically, the promoters of BAX and PPM1D were primarily regulated by *f*, whereas MDM2 and CDKN1A exhibited additional regulation through *μ*. Following IR, promoter regulation patterns resembled those observed in Nutlin-treated cells, despite the different p53 accumulation dynamics (**Figure 1B**). Upon etoposide treatment, CDKN1A promoter remained predominantly regulated by *f*. BAX also followed this pattern, though with a reduced magnitude of *f* and a different temporal profile. In contrast, MDM2 and PPM1D promoters showed increased regulation through promoter activation and a stronger contribution of *δ*. Notably, no significant differences in regulation were observed between the two etoposide doses. Upon UV radiation, we observed greater variability in promoter regulation across genes and doses. As indicated by the RNA measurements, most promoters exhibited attenuated activity despite high p53 levels (**Figure 2E-F**), likely due to the UV-induced decrease in transcription (**Supplementary Figure 4**). At low UV doses, BAX, MDM2 and CDKN1A promoters showed minimal regulation across all three parameters, except for MDM2, which displayed slight regulation through *δ.* In contrast, the PPM1D promoter showed an increase in *f*. At high UV doses, PPM1D and BAX clustered together, with *μ* and *δ* playing a major role in their promoter regulation. Meanwhile, MDM2 exhibited remarkably high regulation through *f* across time points. Ultimately, following 5FU treatment, PPM1D and CDKN1A promoters exhibited weak regulation across transcriptional parameters, whereas MDM2 and BAX promoters displayed an *f*-driven regulation of their activity.

Taken together, our results demonstrate that the implementation of our Bayesian inference-based approach enabled the successful identification of differential regulation of transcriptional bursting in p53 target genes in response to diverse stimuli. Consistent with previous studies, most promoters predominantly modulate the frequency of activation and deactivation rates upon induction.

### 4. p53 post-translational modifications contribute to the regulation of promoter activity

Our data unexpectedly showed that gene- and stimulus-specific promoter activity is uncoupled from p53 dynamics. We hypothesized that this could be due to differential binding of p53 to its response elements (REs). Therefore, we performed Chromatin Immunoprecipitation (ChIP) experiments to quantify the fraction of p53 bound to REs at gene promoters. Strong p53 binding was observed following IR and chemotherapeutic treatments, with no substantial differences between these conditions. Interestingly, low UV doses resulted in p53 binding levels comparable to basal conditions, while higher UV doses led to increased binding (**Figure 4A**). Surprisingly, these findings showed only a weak correlation with total p53 detected by western blot analysis (**Supplementary Figure 1**). As p53 binding to its target gene promoters was not able to explain the observed gene- and stimulus-dependent promoter activity, we explored further regulatory mechanism. Many studies reported that post-translational modifications at the p53 C-terminal domain (CTD) play a critical role in transcriptional regulation. Particularly, acetylation at the residues K370 and K382 is associated with increased target gene expression (**Figure 4B**). *In vitro* and *in vivo* studies have demonstrated that acetylated p53 is enriched at the CDKN1A promoter compared to total p53 (37). To compare total and acetylated p53 levels across different stimuli, we performed western blot analysis on A549 cells harvested at selected time points corresponding to peak p53 accumulation. Consistent with time-lapse microscopy observations, p53 showed different pattern of accumulation depending on the applied stimulus. Notably, despite high total p53 levels under certain conditions, its acetylation state was unexpectedly low in many cases, as exemplified by etoposide, 5-FU and low UV treatments (**Figure 4C**). Importantly, p53 acetylation mostly correlated with smFISH measurements of RNA levels. For example, at the 3-hour time point, IR and Nutlin treatment led to higher expression of target genes compared to etoposide and 5FU (**Figure 2D-J**).

**Figure 4.**
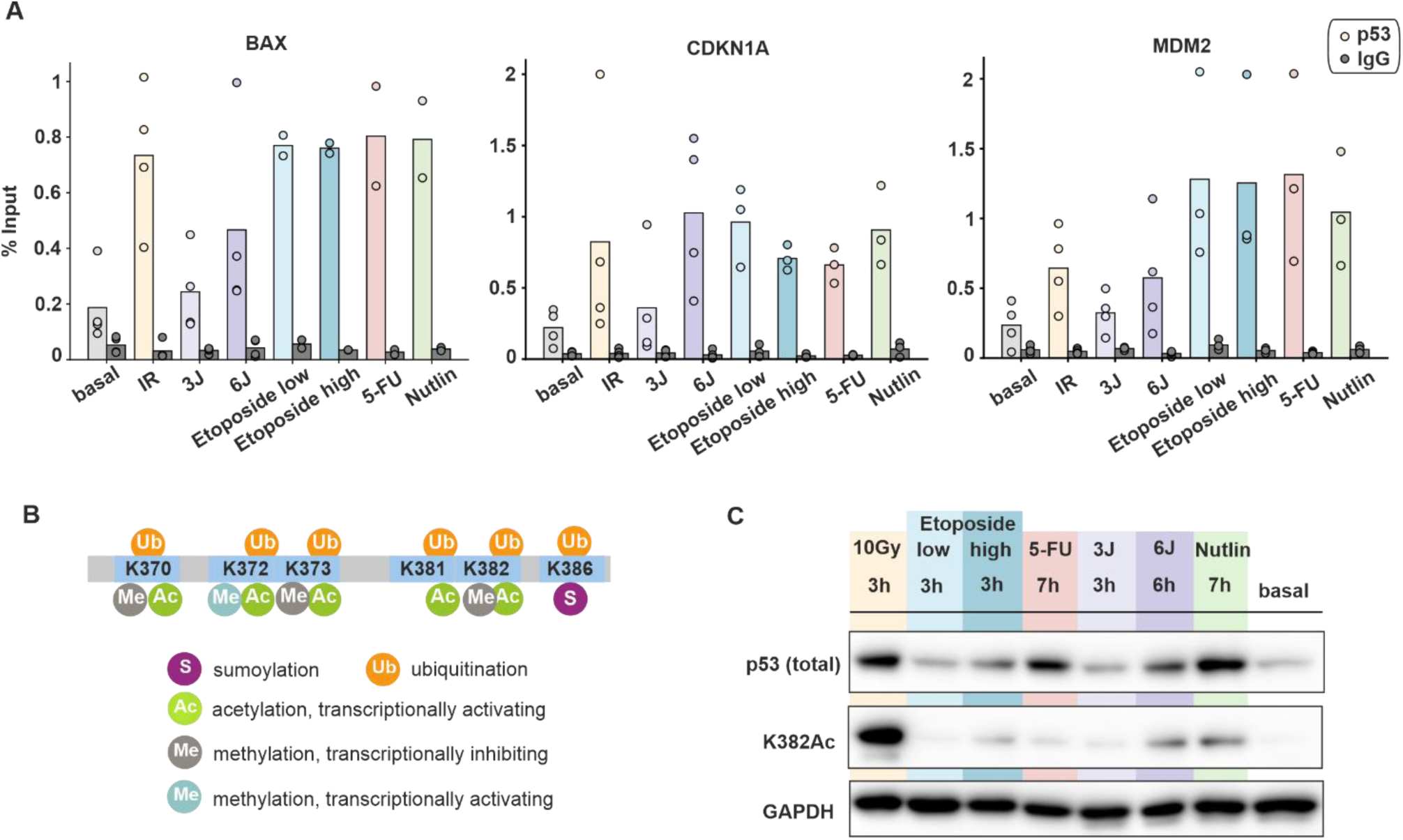
Post-translational modifications and promoter binding of p53 are uncoupled from its dynamics. (**A**) Amount of p53 bound to indicated target gene promoters upon different stimuli measured by ChIP and calculated as percentage of input. Individual data points (mean values of triplicate quantification in RT-qPCR measurements) from three to four biological repeats are shown as dots; mean values are displayed as bars. (**B**) Schematic illustration of p53’s C-terminal modifications (modified from Friedrich *et al.* 2019). (**C**) Comparison of total p53 and acetylated p53 at K382 upon different stimuli through Western Blot assays. The time points were chosen based on the peak p53 levels measured in population assays. GAPDH was used as loading control. Data is representative of three independent repeats.

Taken together, our results indicate that total p53 levels alone do not predict transcriptional activity. Instead, acetylation of the p53 CTD appears to be implicated in determining gene activation. This suggests that post-translational modifications act as a regulatory layer that fine-tunes p53-dependent transcriptional responses to different stimuli. Overall, we provided evidence that p53-mediated gene expression is not solely dictated by its accumulation or DNA binding, but rather by the integration of multiple regulatory mechanisms.

## Discussion

In this study, we systematically compared the dynamic p53 response and the induction of its target genes upon multiple sources of genotoxic stress. Distinct p53 dynamics enable cells to respond differently to various stimuli, ultimately influencing cell fate. For example, pulsatile p53 primarily induces the expression of genes involved in transient responses to DNA damage (cell cycle arrest), while sustained p53 is more closely associated with terminal outcomes (apoptosis, senescence) (38). Therefore, understanding how different p53 dynamics regulate gene expression is crucial for understanding its role in tumor suppression. Our data reveal that the dynamics of p53 do not always correlate with the corresponding gene expression, indicating an uncoupling of TF levels and the activity of its target gene promoters. This disconnect has been observed in other studies (21) and for other TFs, such as SMAD2/4 where TGF-β-dependent nuclear translocation does not align with the transcription profile of its target gene *ctgf* (39). Although TFs are key players in gene expression, they are not the sole determinants.

In our study, we induced various types of DNA damage that could directly interfere with transcription. For instance, following UV radiation, we observed a general reduction in newly synthesized RNA. This is consistent with previous findings, showing that UV radiation not only inhibits transcriptional initiation, but also slows transcript elongation due to Pol II stalling (36,40,41). We hypothesize that in these cases, p53 acts as a sensor of DNA lesions: it accumulates upon UV exposure but can only activate its gene expression response once the damage is repaired by the nucleotide excision repair machinery. This is evidenced by the increase in mRNA levels for most target genes only at later time points and the fact that promoter activity is not strongly regulated by the transcription rate following UV exposure. Similarly, 5-FU treatment also stabilizes p53 by disrupting the MDM2-p53 feedback circuit (34). 5-FU misincorporation is known to disrupt many aspects of RNA metabolism, particularly affecting rRNA, tRNA and snRNA (42,43). However, its potential impact on mRNA processing remains unclear.

To build a comprehensive dataset for studying stochastic gene expression, we employed smFISH. This powerful tool offers single-molecule resolution and spatial information within individual cells. It provides high sensitivity and specificity, avoids amplification biases and enables multiplexing to study several genes simultaneously (44). However, it is limited by low temporal resolution in fixed cells and the ability to analyze only a relatively small number of cells. Therefore, researcher have employed live-cell imaging of transcriptional dynamics to follow cells over time (18,45) and be able to capture the timescale of transcription, which ranges from minutes to hours in mammalian cells. Neverthless, dual-color smFISH allowed us to gain insights into transcriptional bursting. By fitting a telegraph model to our smFISH data using Bayesian inference, we calibrated five model parameters and systematically characterized features of promoter activity in a stimulus-specific manner. Consistent with other studies (21,24), our results reveal that p53-dependent transcription is mainly regulated by the activation and deactivation rates of the promoter. Although p53 accumulation drives promoter activation, its temporally changing levels do not directly lead to differential promoter regulation. For example, increasing p53 levels following Nutlin treatment and pulsatile p53 dynamics upon IR result in similar regulation of *f*, *μ* and *δ*. Moreover, gene expression appears to be unaffected by p53 concentration. For instance, despite etoposide inducing vastly different p53 levels at different concentrations, the response profiles of target genes remained largely similar.

The telegraph model, i.e. the two-state promoter model with Markovian switching, is a simple, accessible model that often represents bursting kinetics of gene transcription sufficiently well and is thus commonly used (11). In our application, however, measured fluorescence from nascent RNA at active TSs was overdispersed w.r.t the model prediction. We compensated this effect by the addition of a phenomenological variance parameter, but a mechanistic description is lacking. While the base model itself stayed unaltered, we derived an explicit expression for the distribution of the total RNA count at steady state in the presence of multiple TSs (telegraph models with identical parameters, indistinguishable from data) that can be applied to polyploid cells, and for which we could show good agreement between model fit and data. Bayesian inference provided us with two important advantages compared to frequentist point estimates (e.g. taking only the most likely combination of parameters). First, the posterior distribution quantifies the remaining uncertainty in the model parameters after training on the experimental data in a principled way. Second, it gave us access to the minimum-mean-square-error (MMSE) estimator for the parameters which has many desirable statistical properties (46). While MCMC is a versatile tool to implement Bayesian inference, it requires a fair amount of expert knowledge to adjust in order to achieve its desired statistical properties. At the same time, single-chain MCMC can perform poorly in scenarios with multiple distant posterior modes, often unknowingly so to the user (21). Finally, Bayesian inference with MCMC comes with a high computational cost, though this is often outweighed by the expense of generating experimental data in the lab.

Moreover, as model parameters of CDKN1A show some inconsistencies among biological replicates, some interpretation should be drawn with caution. However, it is important to highlight that stochastic gene expression leads by itself to heterogeneity. Most of the genes are transcribed in bursts at low and high expression levels. However higher RNA levels are mainly dominated by extrinsic noise (24,47). Additionally, TF dynamics positively affect noise in gene expression.

Although the random telegraph model could help us describing how the transcriptional parameters change in response to varying stimuli, our findings suggest that underlying levels of promoter regulation are not fully captured by a two-state promoter model. For instance, more complex models incorporate feedback regulation, multiple gene activation steps, and multiple gene activation pathways (11). Recent studies have integrated transcriptional bursting with genomic architecture and chromatin accessibility (48), revealing that these factors preferentially affect burst frequency (BF). Using high-throughput imaging screens, researchers demonstrated that histone acetylation influences BF and TS intensity, primarily by reducing off-times (49). Therefore, profiling histone modifications at p53 target genes could help unravel promoter regulation. However, bioinformatic databases provide only partial profiles of acetylation marks, such as the canonical H3K27Ac and H3K9me3. Additionally, most analysis are performed in untreated cells or following Nutlin/DMSO treatments (50). Notably, DNA damage induces numerous post-translational modifications not only of the TF, leading to a stabilization of p53 levels in the nucleus, but also affecting histones and other co-factors. In future studies it will be interesting to implement histone modifications of p53 target gene promoters to investigate features of transcriptional bursting in more detail.

The interaction between chromatin and TF is relatively brief, and p53 residence time is modulated by C-terminal acetylation upon DNA damage induction (51). Recent studies have observed changes in CTD acetylation between the first and the 2nd p53 pulses upon IR, which may explain the regulation of target genes with oscillatory expression. In our study, we expanded the investigation of CTD PTMs to multiple sources of both p53 induction and DNA damage. We observed significant differences across stimuli, with the highest levels of p53 acetylation occurring after IR and Nutlin treatment. Interestingly, these conditions also exhibit similar *f*-driven promoter regulation. Moreover, total p53 levels do not correlate with its acetylation state, suggesting that PTMs fine-tune promoter activation. Indeed, p53 acetylation may enhance transcriptional regulation by increasing site-specific DNA binding activity (37). However, our findings show that p53 binding at the promoter remains consistent across target genes and treatments, except for low doses of UV. These results align with previous studies showing conserved p53 DNA binding across treatments and cell-types (52). It has been recently shown that spatio-temporal genome organization is directly affected by p53 activation, through its binding to DNA, and indirectly by other factors transcriptionally regulated by p53 (53). Therefore, PTMs and p53 binding alone cannot fully explain the regulation of promoter activity, highlighting the contribution of co-regulatory factors recruited at the site of transcription. Additionally, studies have demonstrated that mRNA half-life plays a crucial role in determining differential target gene expression of pulsatile transcription factors like p53 (54). With our study, we present a systematic analysis of stimulus-dependent stochastic gene expression, setting the stage for further exploration of the complex mechanism controlling p53 target gene expression and ultimately cell fate upon damage induction.

## Data availability statement

All measurements derived from smFISH experiments as well as additional output from Bayesian inference are publicly available via the institutional repository of Technical University Darmstadt (https://doi.org/10.48328/tudatalib-1704). This includes separate plots of the parameters (*f*, µ, δ) for each gene and condition, histogram fits of the active TS exon fluorescence data, histogram fits of the RNA counts, plot of MCMC convergence, posterior distribution of the parameters inference and fitting of TS quantification. Raw image data are available from the corresponding authors upon reasonable request due to their large size. The corresponding analysis code is available at the same link.

## Author contributions

A.L. and F.V. designed experiments and conceived the study. F.V. performed experiments and analysed data. M.S. formulated the mathematical model. N.E. and M.S-A. wrote code and performed the Bayesian inference. F.V. prepared figures with contributions from N.E. F.V., N.E. and A.L. wrote the manuscript, with contributions from all authors. A.L. and H.K. supervised the research.

## Acknowledgments

We thank Petra Snyder (Technical University Darmstadt) for technical assistance, Dhana Friedrich for help in setting up smFISH experiments, and members of the Loewer and Koeppl labs for helpful discussion.

