## Supplementary material for "Decoding stimulus-specific regulation of promoter activity of p53 target genes"

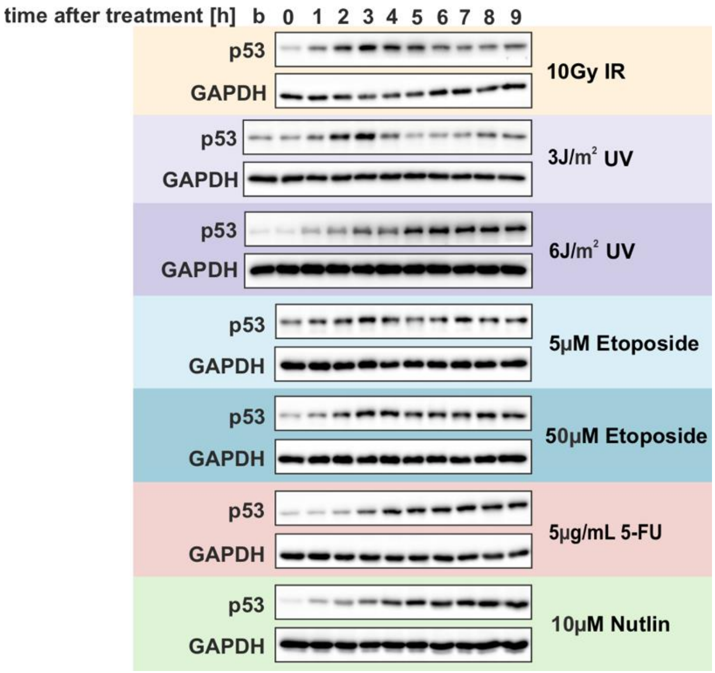


**Supplementary Figure 1 – p53 levels were characterized in A549 wild-type cell line.**

Western Blot measurements of total p53. A549 wt cells were harvested at the indicated time points upon different stimuli. GAPDH was used as loading control. b:basal. Data is representative of three independent repeats.

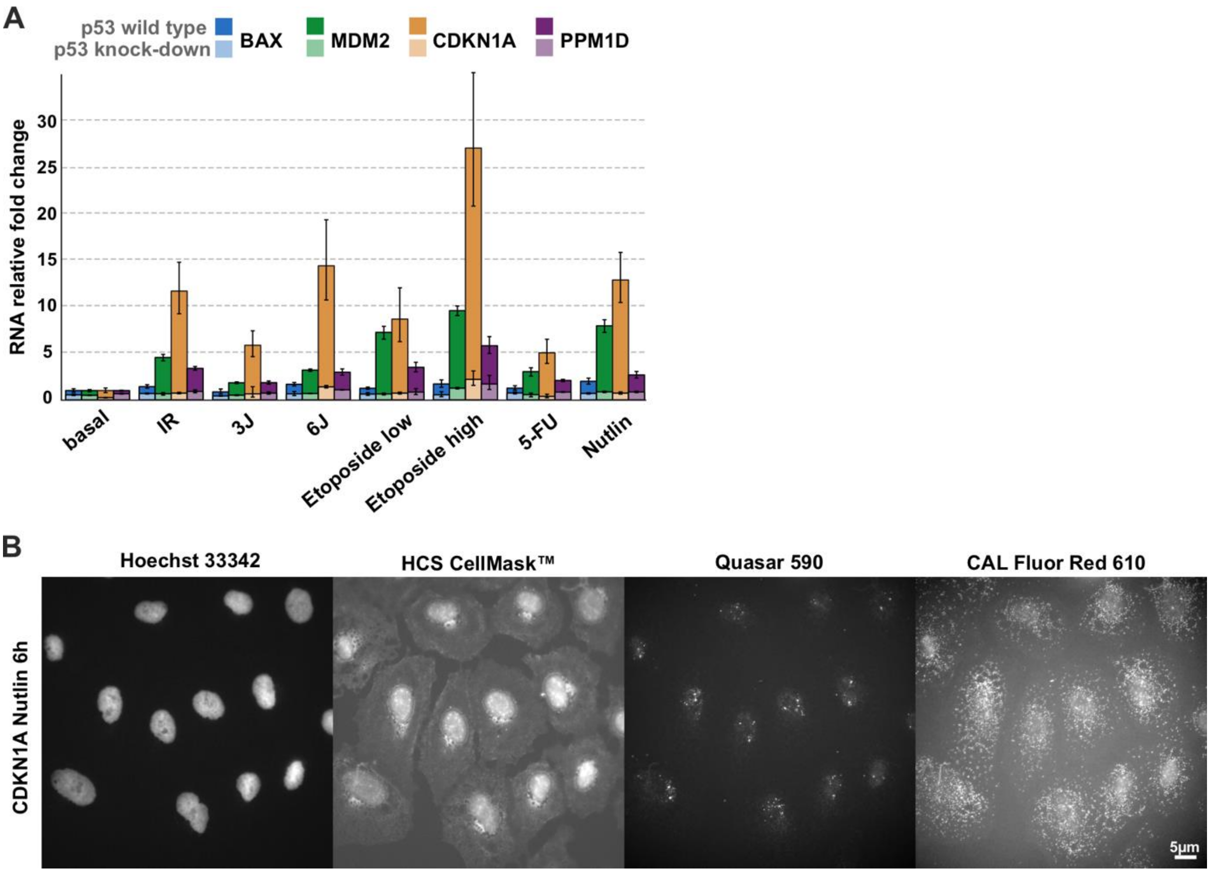


**Supplementary Figure 2A – Expression of p53 target gene in individual cells and cell population.**

**(A)** Expression of selected p53 target genes after treatment in A549 wild-type and p53 knockdown cells. RNA levels were measured by RT–qPCR. The following time points were chosen: 3h post IR, 3J/m$^2$ UV and etoposide low treatments, 6h after 6J/m$^2$ UV and etoposide high treatments, 7h after 5-FU and Nutlin addition. Fold changes relative to basal levels are shown for each cell line as mean and standard deviation from technical triplicates. **(B)** Multicolor fluorescent imaging of exemplary smFISH data. From left to right: individual images of nuclei stained with Hoechst-33342, cytoplasmic staining with HCS CellMask™, exon staining with Cal Fluor 610, and intron staining with Quasar 670 dye. Scale bar corresponds to 5µm distance; images were contrast and brightness enhanced for better visualization.

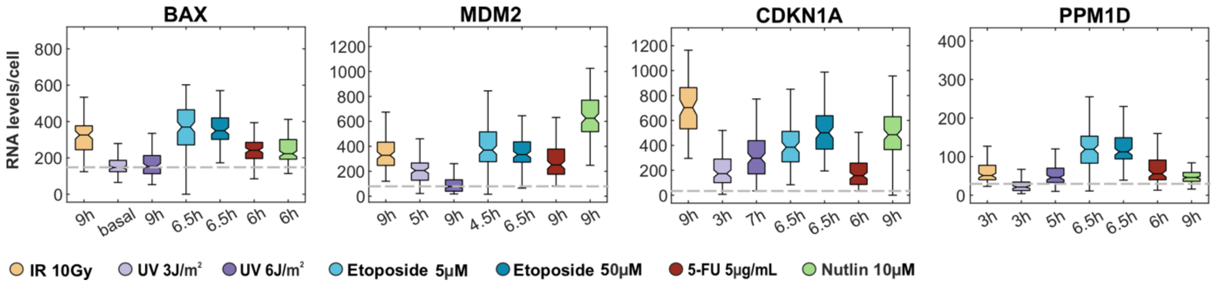


**Supplementary Figure 3 – Comparison of mRNA counts across genes and treatments.**

Quantification of RNA per cell using FISH-Quant for four target genes, presented as box plots. Time points corresponding to peak RNA levels were selected to compare gene expression across different stimuli. Solid lines indicate the mean of each distribution, while boxes represent the interquartile range ($25^{th}$-$75^{th}$ percentiles). Dashed lines denote the mean RNA levels under basal conditions, calculated across all the stimuli for each gene.

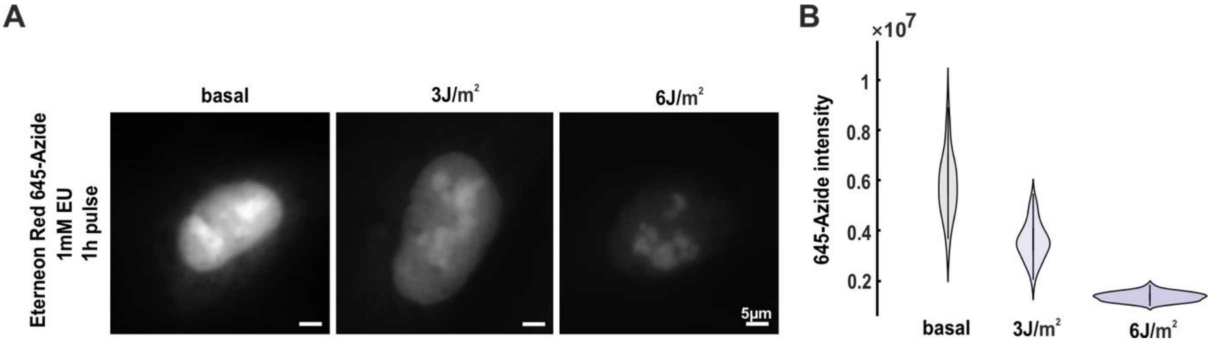


**Supplementary Figure 4 - RNA synthesis is globally inhibited upon UV radiation.**

**(A)** Representative images of metabolic labelling of nascent RNAs, showing decreasing of RNA synthesis after two doses of UV radiations. A549 wt cells were incubated with 1mM EU for 1h and fixed 3h after treatment. They were subsequently stained with 20µM Eterneon Red 645-Azide and Hoechst-33342. **(B)** Fluorescence intensity was then quantified via ImageJ. Scale bar corresponds to 5µm distance; images were contrast and brightness enhanced for better visualization.

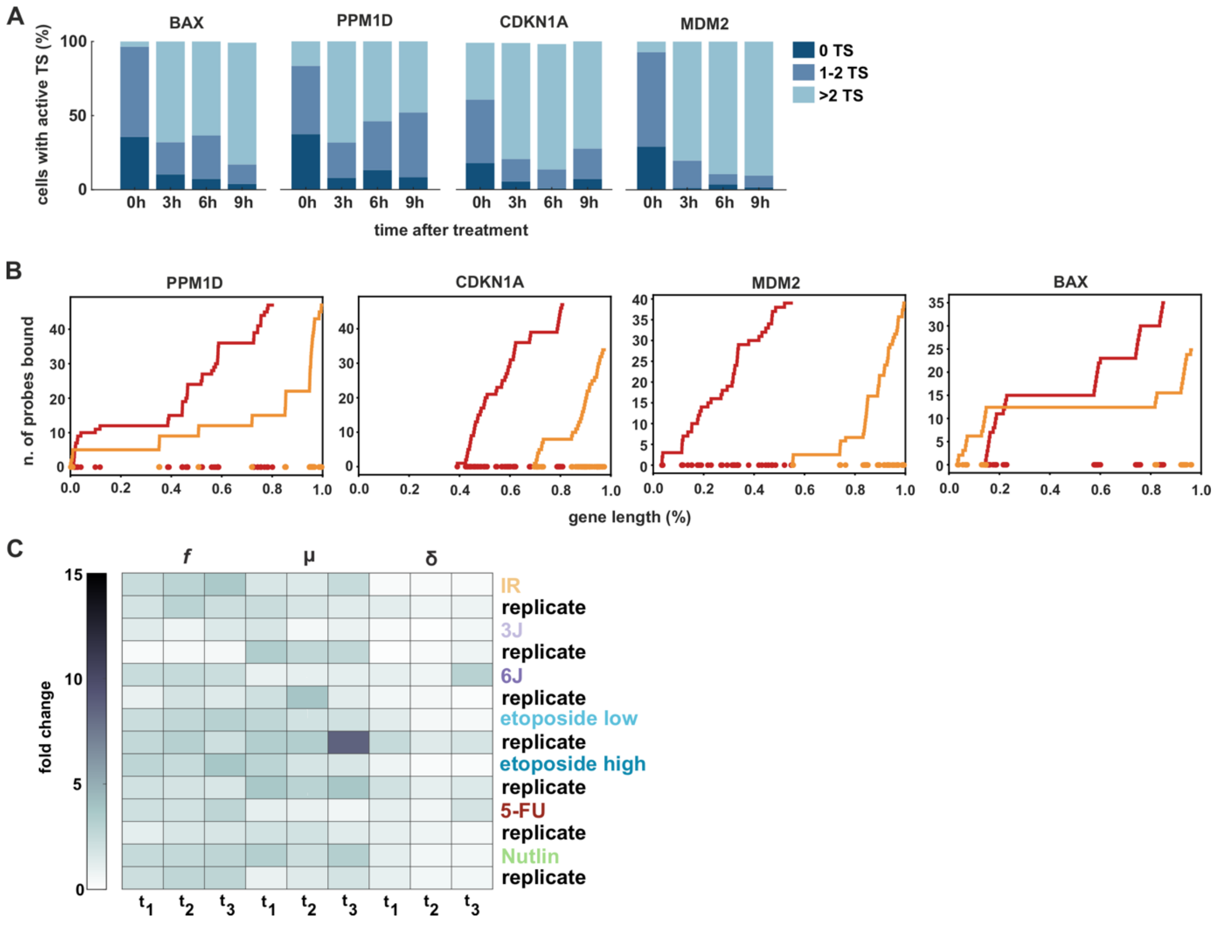


**Supplementary Figure 5A - Analysis of promoter activity for p53 target genes.**

**(A)** Stacked bar plots representing the percentage of cells with 0, 1 or more than 2 active transcription sites (TSs). Data are obtained from smFISH measurements after Nutlin treatment at the selected time points.The maximum number of genomic loci was previously validated in Friedrich et al. 2019 as follows: BAX (4), PPM1D (4), CDKN1A (4), and MDM2 (3). **(B)** Plots show the number of intron (in red) and exon (in yellow) probes bound to transcribing Pol II (y-axis) along the genomic sequence of each p53 target gene, shown in percentage (x-axis). Reference sequences (hg38) were used without 3' and 5' UTRs: chr19:48956199-48961097 (BAX), chr12:68808478-68839849 (MDM2), chr6:36684102-36685800 (CDKN1A), chr17:60600415-60663552 (PPM1D). **(C)** MCMC estimates of f, μ, and δ represented in a heatmap. Each row represents a stimulus, and each column a time point (3h, 6h and 9h for IR, 5-FU and Nutlin; 2.5h, 4.5h and 6.5h for etoposide low and high; 0h, 5h, 7h, 9h for UV 3J/$m^2$ and 6J/$m^2$). Data were normalized to basal conditions and shown as fold changes, representing biological replicates for each condition.
